# The method of modeling the human EEG by calculating radial traveling waves on the folded surface of the human cerebral cortex

**DOI:** 10.1101/242412

**Authors:** Vitaly M. Verkhlyutov, Vladislav V. Balaev

## Abstract

There are many data about traveling waves in the cortex of animals such as rats, ferrets, monkey, and even birds. Waves registered invasively using electrical and optical imaging techniques. Such registration is not possible in healthy man.

Non-invasive EEG recordings show scalp waves propagation at rates two orders greater than the data obtained invasively in animal experiments. At the same time, it has recently been argued that the traveling waves of both local and global nature do exist in the human cortex. We have developed a novel methodology for simulation of EEG as produced by depolarization waves with parameters taken from animal models. We simulate radially propagating waves, taking into account the complex geometry of the surface of the gyri and sulci in the areas of the visual, motor, somatosensory and auditory cortex. The dynamics of the distribution of electrical fields on the scalp in our simulations is consistent with the EEG data recorded in humans.

## Introduction

Traveling waves (TW) on the cerebral cortex surface were discovered in the 30s of the 20th century (Adrian & Mathews, 1934), and were then detected in human via the EEG mapping (Adrian & Yamagiva, 1935; Lindsley, 1938) and then were investigated minutely in 50–60s (Walter, 1953; Petsche, 1955; Shipton, 1957; Anan’iev et al., 1956; Livanov & Anan’iev, 1960, Monakhov, 1961; Dubikaitis and Dubikaitis, 1960, Raimond, 1961) and less intensively in subsequent years (Hughes, 1995). At present, interest in this phenomenon returned for reasons the registration techniques improvement (Ferrea, 2012) and owing to a number of physiological discoveries (Biswal et al., 1995; Hindriks et al., 2014; Matsui et al., 2016), which may explain of the TW.

TW on the surface of the scalp have a speed of about 5–15 m/s (Alexander et al., 2013), while lower speed was recorded in electrocorticography (ECoG) experiments (Zhang et al., 2017). Moreover, when recorded using microelectrode matrices placed on the cortical surface, the velocity of the TW appeared lower than 0.5 m/s (Martinet et al., 2017) which is similar to the speed detected in animals by means of electrical (Rubino et al., 2006; Ferezou et al., 2007; Reimer et al. al., 2011; Takahashi et al., 2011) or optical recordings (Muller et al., 2014). Importantly, when registered within a small area about 16 mm^2^, simpler TW configurations appear, which in many cases might be interpreted as radial waves propagating from one epicenter (Martinet et al., 2017).

Previously, we proposed an elementary model for the TW spread within the human visual cortex, which explained all known to us at that time phenomena of EEG TW in alpha-range on the head (Verkhliutov, 1996). However, the primitiveness of this model made it necessary to search ways of calculating TW in a more realistic model. This became possible when the appearance of the Boundary Element Method (BEM) based on the high quality MRI images (Fuchs et al., 1998) and algorithms for geodetic distances calculation on complex surfaces (Siek et al. 2002).

In this report we propose an EEG data generatin method consistent with the intracortical hypothesis (Hindriks, 2014), which suggests that the main activity recorded in the EEG is related to the TWs of electrical potentials propagated by intra-cortical short fibers (Ermentrout & Kleinfeld, 2001; Han et al., 2008; Zheng & Yao, 2012). We limit our attention to only radially propagating omni-directional waves in the 10 Hz range.

We also assume that spontaneous TWs are not a constant process on the cortex surface and we believe that TW are more related to rest, sleep, pathological and epileptic activity (Wu et al., 2008). The spontaneous TWs are suppressed during sensory stimulation (Muller et al., 2014). At the same time the evoked and motor potentials exhibit TW-like nature (Rubino et al., 2006).

## Results

Dynamic distributions of dipoles (Fig.2C) from concentrically propagating potential waves (Fig 2D, 3C) from 28 epicenters located at various cortical points (Fig. 1) were calculated. We simulated 28 128-channel EEG tracks (Fig.2A, 3A & Supplementary materials) and calculated the distribution of EEG potentials on the scalp (Fig. 2B; 3B & Supplementary materials). In all of the cases the standard localization methods did not manage to reproduce the wave-like potential distribution on the cortical surface (Fig. 3D & Supplementary materials). We also reproduced traveling waves on the scalp (see Supplementary materials).

**Figure 1:**
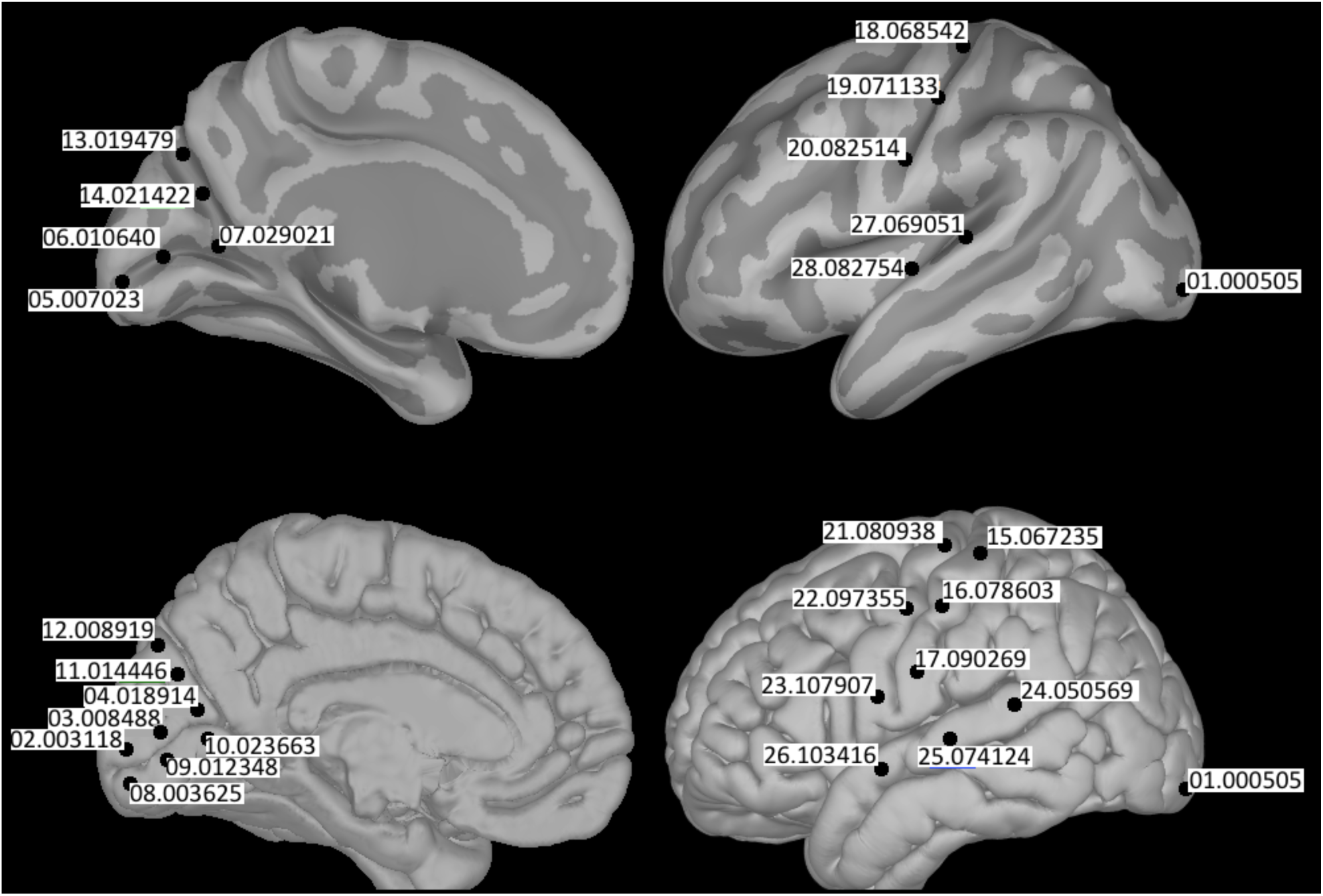
Epicenters for model waves. The epicenters are indicated by black circles and correspond to the order numbers and to the indices of the vertices of the left hemisphere of the cortical surface ICBM152 from Brainstorm data base (Tadel et al., 2011). Sagittal and lateral inflated surfaces - at the top, sagittal and lateral no inflated surfaces - at the bottom.

**Figure 2:**
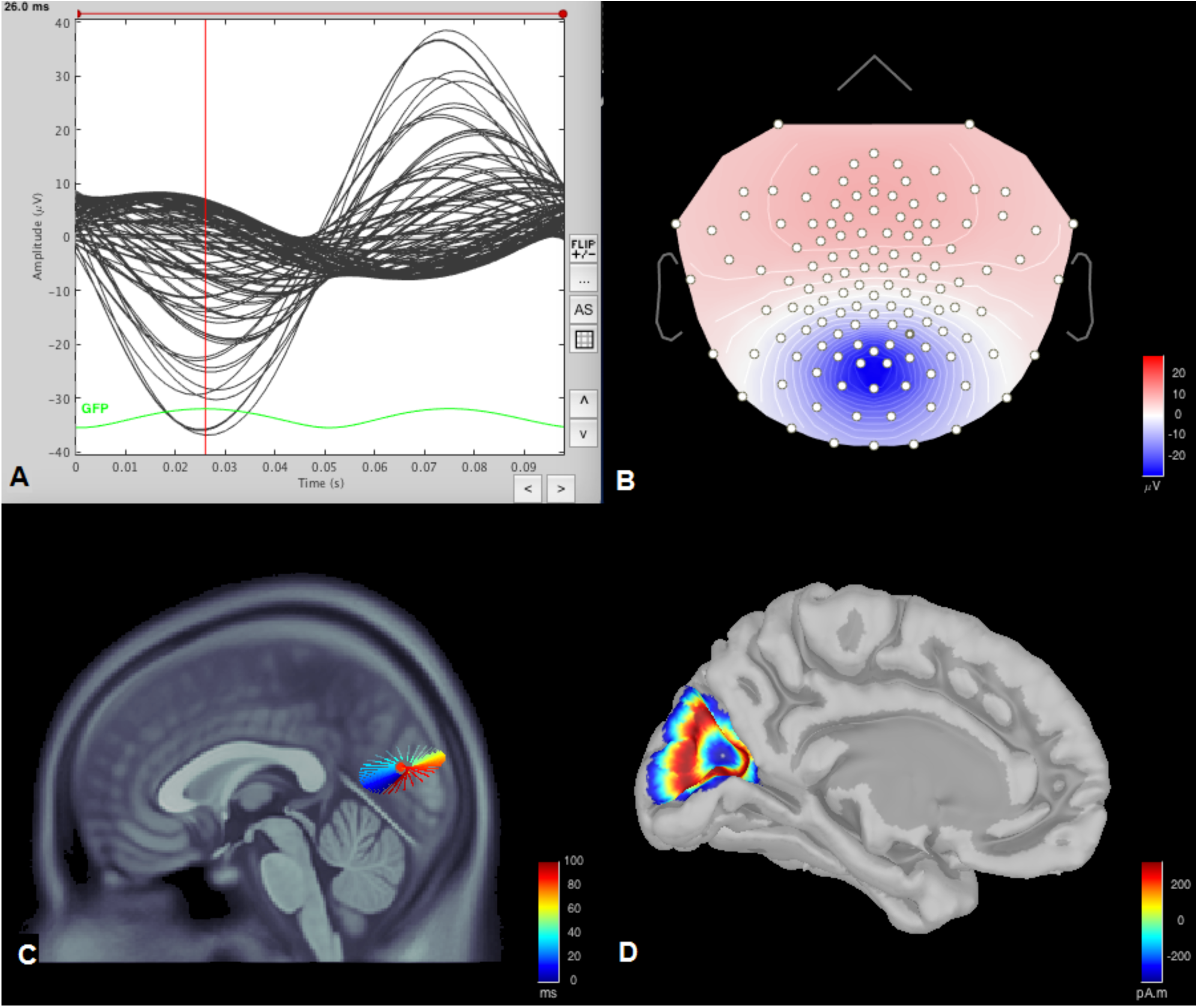
Modeling the EEG from a radial wave in the visual cortex. (A) 100 ms of simulated EEG, (B) electric field distribution on the “scalp” from two symmetric waves in both hemispheres with an epicenter at the vertex 04.18914 of the cortical surface model ICBN152 (the second wave is mirrored with respect to the sagittal plane), (C) the equivalent dipole corresponding to the time moment marked by the cursor in (A), (D) the current density distribution from model radial wave on the cortical surface.

**Figure 3:**
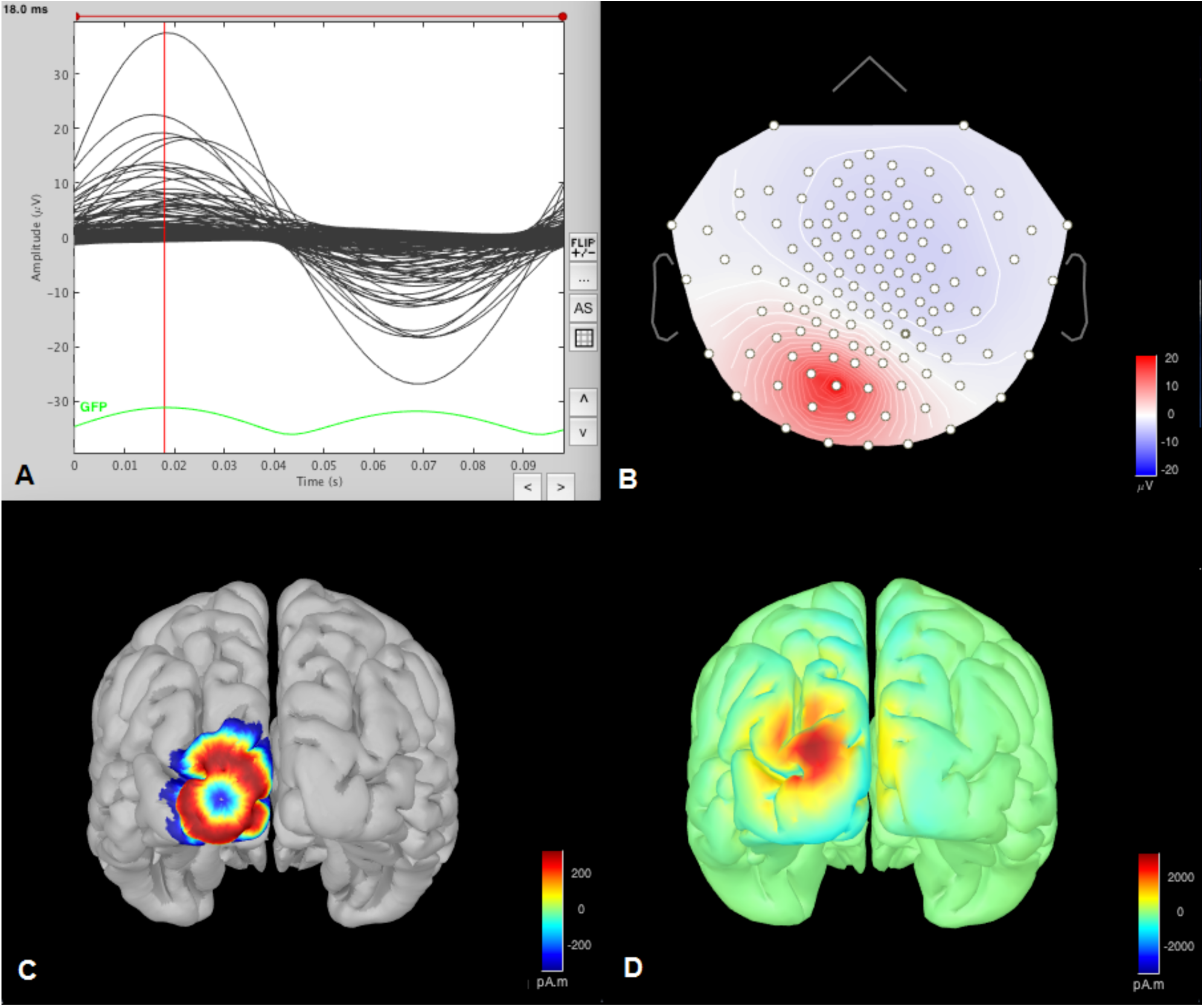
Modeled EEG from the wave propagation of potentials with the epicenter at the occipital pole. (A) Modeled EEG for 100 ms. (B) Electric field distribution on the “scalp” from wave in left hemisphere with an epicenter at the vertex 01.000505 of the cortical surface model ICBN152. (C) The current density distribution from radial wave on the cortical surface model. (D) The current density distribution on the “cortex” localized by the wMNE standard method.

Table 1 summarizes the data: the epicenters of modeled waves, the anatomical structures their pertain to, the functional cortical networks in which the wave process is assumed to propagate, the EEG rhythms, the maximum given current density in the cortex, the maximum potential amplitude on the scalp surface, dynamics of the distribution of potentials on the surface of the scalp.

**Table 1.** The maximum amplitude of the modeled EEG and the type of moving waves on the scalp from radial waves propagating from the epicenters located in the visual, sensorimotor and auditory cortex of the human brain’s ICBN152 model.

| № | Vertex | MNI | Structure | RSN | Rhythm | CD | SPL | STWL | SPB | STWB |
| --- | --- | --- | --- | --- | --- | --- | --- | --- | --- | --- |
| 1 | 01.000505 | -18.8 -102.7 -3.2 | Pol.occipital. | VIS | alpha | ±50 | ±40 | RL | ±50 | 2OF |
| 2 | 04.018914 | -0.0 -73.3 15.3 | Cuneus | VIS | alpha | ±50 | ±20 | RR | ±40 | FO |
| 3 | 03.008488 | 0.5 -84.4 7.3 | Cuneus | VIS | alpha | ±50 | ±25 | L | ±45 | OF |
| 4 | 02.003118 | 0.6 -93.8 2.6 | Cuneus | VIS | alpha | ±50 | ±40 | RR | ±80 | FO |
| 5 | 11.014446 | 0.4 -76.9 24.4 | Cuneus | VIS | alpha | ±50 | ±45 | RR | ±80 | FO |
| 6 | 12.008919 | -0.0 -83.2 34.1 | Cuneus | VIS | alpha | ±50 | ±15 | RR | ±30 | FO |
| 7 | 10.023663 | -0.9 -66.8 3.0 | G.lingualis | VIS | alpha | ±50 | ±20 | RA | ±30 | OF |
| 8 | 09.012348 | -0.0 -80.2 -2.1 | G.occ.temp. | VIS | alpha | ±50 | ±20 | R | ±35 | OF |
| 9 | 08.003625 | -0.6 -93.0 -10.7 | G.occ.temp. | VIS | alpha | ±50 | ±20 | OF | ±35 | OF |
| 10 | 07.029021 | -28.1 -62.8 5.7 | S.calcarinus | VIS | alpha | ±50 | ±7 | RP | ±9 | 2FO |
| 11 | 06.010640 | -12.3 -80.5 4.2 | S.calcarinus | VIS | alpha | ±50 | ±20 | L | ±7 | FO |
| 12 | 05.007023 | -4.5 -87.7 -0.8 | S.calcarinus | VIS | alpha | ±50 | ±15 | RR | ±30 | OF |
| 13 | 14.021422 | -15.4 -68.5 26.5 | S.p.occip. | VIS | alpha | ±50 | ±20 | RP | ±40 | FO |
| 14 | 13.019479 | -14.6 -71.3 34.7 | S.p.occip. | VIS | alpha | ±50 | ±20 | RP | ±35 | FO |
| 15 | 15.067235 | -47.6 -28.9 64.0 | G.postcentr. | SSM | mu | ±50 | ±15 | RL | ±15 | 2FO |
| 16 | 16.078603 | -59.1 -18.8 49.3 | G.postcentr. | SSM | mu | ±50 | ±15 | LP | ±25 | 2FO |
| 17 | 17.090269 | -65.6 -9.9 25.3 | G.postcentr. | SSM | mu | ±50 | ±25 | L | ±30 | 2OF |
| 18 | 21.080938 | -37.8 -17.3 69.7 | G.precentr. | SSM | mu | ±50 | ±15 | RP | ±25 | FO |
| 19 | 22.097355 | -56.2 -4.6 47.6 | G.precentr. | SSM | mu | ±50 | ±25 | RR | ±25 | 2OF |
| 20 | 23.107907 | -62.4 4.3 20.3 | G.precentr. | SSM | mu | ±50 | ±20 | RL | ±20 | 2OF |
| 21 | 18.068542 | -22.0 -27.4 55.1 | S.Rolandic | SSM | mu | ±50 | ±30 | RP | ±55 | 2FO |
| 22 | 19.071133 | -34.4 -25.7 49.3 | S.Rolandic | SSM | mu | ±50 | ±30 | FO | ±50 | FO |
| 23 | 20.082514 | -41.8 -15.7 33.7 | S.Rolandic | SSM | mu | ±50 | ±40 | RA | ±30 | 2OF |
| 24 | 24.050569 | -67.1 -41.3 14.8 | G.temp.sup. | AUD | tau | ±50 | ±15 | R | ±20 | 2OF |
| 25 | 25.074124 | -67.9 -23.1 4.9 | G.temp.sup. | AUD | tau | ±50 | ±15 | RR | ±25 | 2FO |
| 26 | 26.103416 | -61.0 0.6 -7.1 | G.temp.sup. | AUD | tau | ±50 | ±25 | RR | ±30 | 2FO |
| 27 | 27.069051 | -61.0 0.6 -7.1 | F.Sylvian | AUD | tau | ±50 | ±25 | L | ±40 | 2OF |
| 28 | 28.082754 | -51.3 -17.0 9.9 | F.Sylvian | AUD | tau | ±50 | ±40 | RL | ±45 | 2OF |

№ п/п, **Vertex**- index of the vertex in the ICBN152 model of the cerebral cortex coinciding with the epicenter of the propagating cortical wave, **MNI** epicenter coordinates in mm, **Structure** – anatomical structures of the epicenter location, **RSN** – functional (resting) brain network, where the epicenter is placed (VIS - visual, SSM - sensorimotor, VIS - auditory), **Rhythm** – assumed rhythm **10 Hz**, **CD** – predetermined maximal current density in the cortex in nA/mm<sup>2</sup>, **SPL**-estimated maximum amplitude of potentials on the scalp in µV of the model EEG in the case of one wave in the left hemisphere, **STWL** - dynamics of the wave on the scalp in the case of one wave in the left hemisphere (OF- occipito-frontal, RL- left rotation,, RR – right rotation, R-right, L-left, RA-right forward (anterior), RP-right backward (posterior), LP-left backward (posterior), CO-from center to occipital lobe), **SPB**- maximum amplitude of potentials on the scalp in µV-scale modeled EEG for symmetric propagating waves in both hemispheres, **STWB**-wave dynamics on the scalp in the case of symmetric propagating waves in both hemispheres (2-bifurkated extrema, FO-fronto-occipital, OF- occipito-frontal, CO-from center to occipital).

The models of alpha rhythm and alpha-like rhythms differed in localization only. The epicenters for the alpha rhythm were placed in the regions of the calcarinus and parieto-occipital sulcuses, the ones for the mu-rhythm placed in the area of the central sulcus, and for the tau rhythm, the epicenters were set in the vicinity of the Silvian fissure.

With an equal wave source current density of ± 50 nA / mm2 in the cortex, the potential amplitudes on the scalp were greatest for the alpha rhythm reaching 160 μV from peak to peak. The same value for the mu-rhythm was 110 μV, and 90 μV for the tau rhythm. Such values agree with those observed experimentally for these rhythms (Markand., 1990).

The most frequent dynamics is TW from frontal to occipital lobe - 10 epicenters out of 14 are observed for alpha sources. The total amplitude in this case was 802 μV. For TW from the occipital to the frontal regions - 290 μV.

TWs from the frontal to the occipital lobe and in the opposite direction for mu-and tau-rhythms are approximately equal for different epicenters. In most cases, for Rolandic and Temporal rhythms, a bifurcation of the field extrem on the scalp surface is observed for symmetrically placed sources in both hemispheres.

The rotation of the patterns in the visual cortex is predominantly clockwise for the source in the left hemisphere. Left rotation is observed for the epicenter at the pole of the occipital lobe. In the other cases, transverse and diagonal TWs are detected. Rotational, diagonal and transverse wave movements are observed in case of an asymmetric source location for mu and tau rhythms without noticeable predominance for any type.

The predominance of the TW movements from the frontal to the occipital regions in the simulation coincides with the experimental occurrence frequency of such patterns for EEG (Fig. 4B).

**Figure 4:**
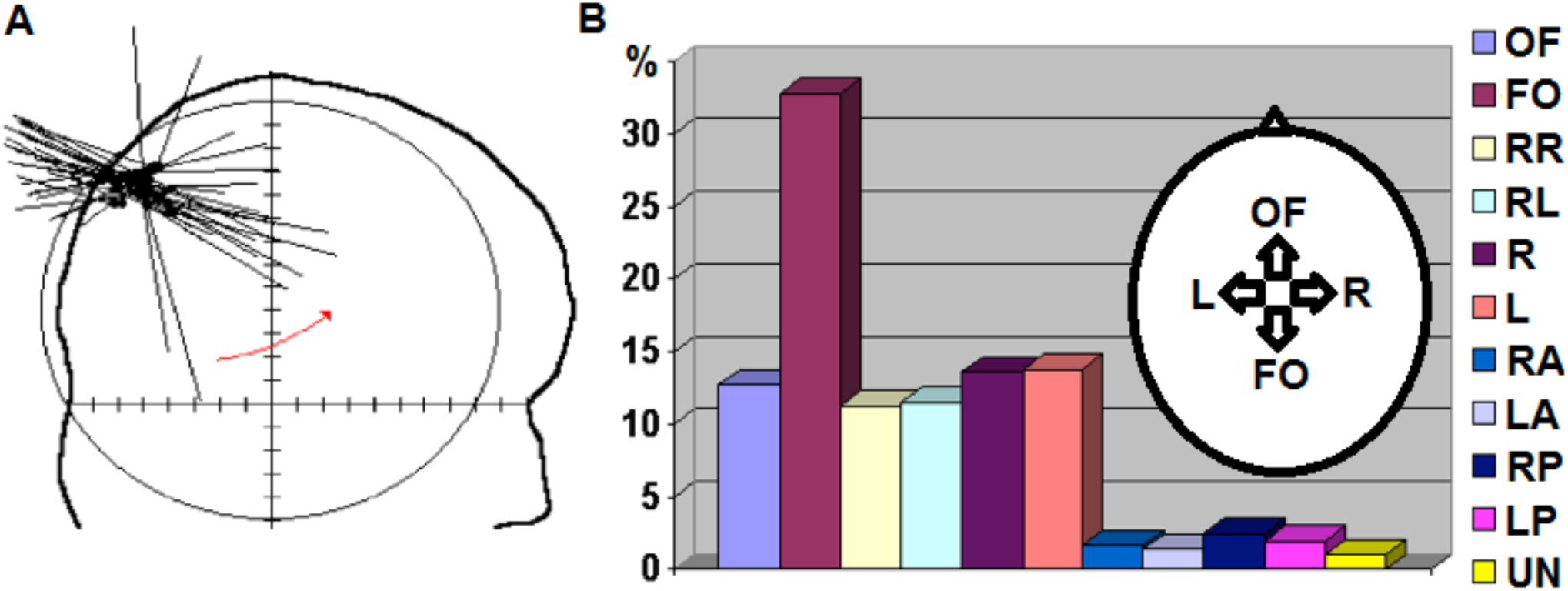
EEG traveling waves. (A) Dynamics of an equivalent dipole in the occipital lobe of the human brain. (B) A diagram illustrating the predominance of frontal-occipital EEG TW on a scalp in an experiment. Data were obtained from 12 healthy subjects at rest with closed eyes (Verkhlyutov, 1999). Types of TW on the scalp: FO-fronto-occipital, OF-occipito-frontal, RR-right rotation, RL-left rotation, R-right movement, L-left movement, RA-right-anterior, LA-left-arterior, RP-right-posterior, LP-left-posterior, UN-unknow.

Since in our model the form of the EEG distribution dynamics on the scalp is related to the location of the epicenters and their symmetry in the hemispheres, the important objective is to recognize such patterns and relate them to the localization of the wave process.

To demonstrate the degree of similarity / difference in the dynamics of 128-channel EEG patterns, we used the corr2 procedure from the MATLAB standard library, which is intended for raster images analysis. The dynamics of the simulated EEG patterns were compared from 28 points in asymmetric and symmetric cases. For both asymmetric and symmetric cases, the average correlation values were below 0.08 and did not differ statistically except for epicenter No. 6. The maximum correlation values for most epicenters were close to 1 (Fig. 5).

**Figure 5:**
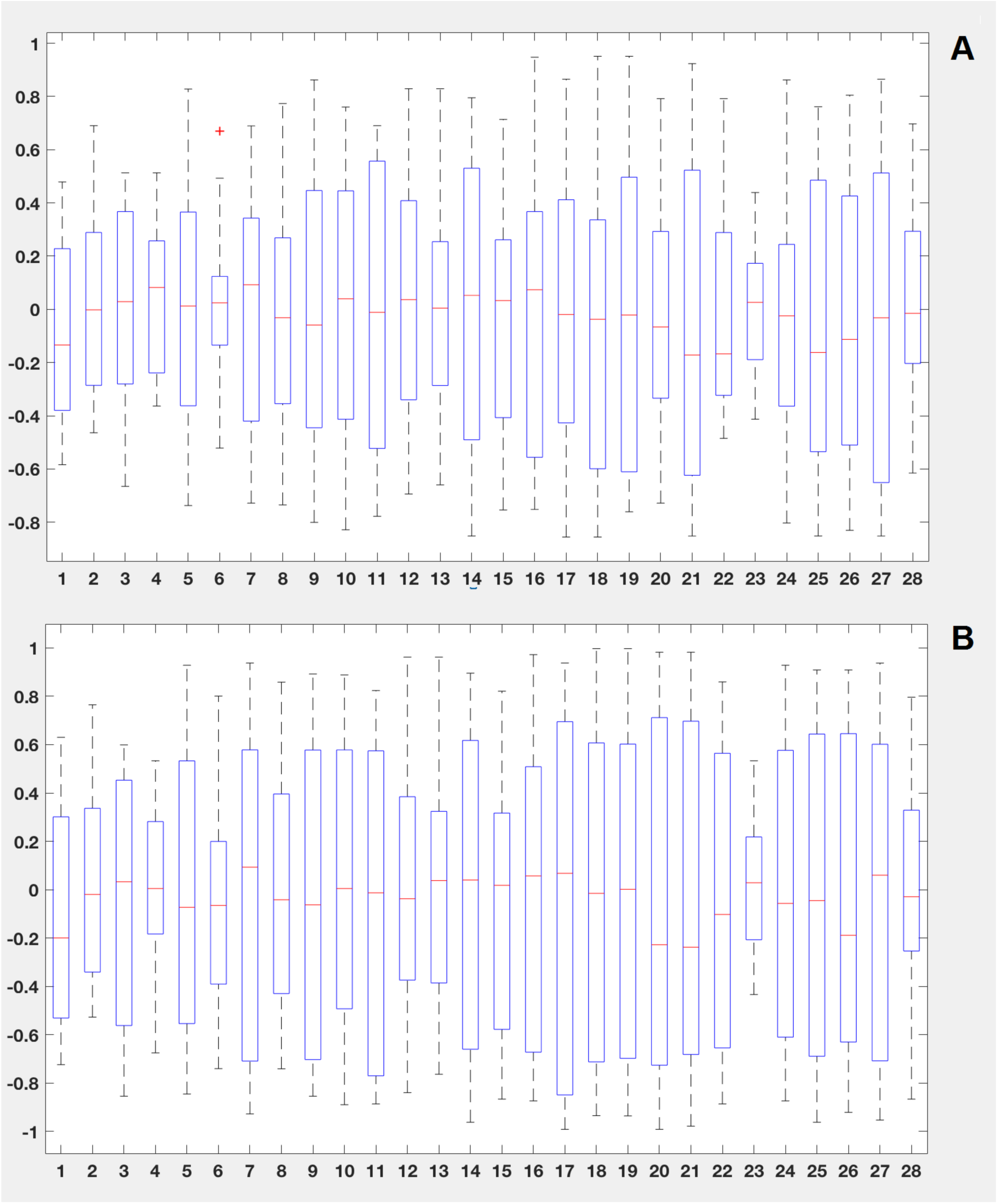
Comparison of mean, minimal and maximal values, first and third quartiles of the correlation of dynamic patterns of simulated EEGs from radial waves with epicenters 1.-28. (Fig. 1) when compared to each other for the left hemisphere (A) and for both hemispheres (B).

## Discussion

According to our model, the dynamics of EEG potentials can point out the epicenter of wave propagation. While modeling, we found that the types of the waves movement occur most often in the direction from the frontal to the occipital regions (FO) and this predominance coincides with the experimental data (Verkhlyutov, 1999; Patten et al., 2012) (Fig. 4). It has also been shown experimentally that the TW from the frontal to the occipital lobe is accompanied by the rotation of an equivalent dipole in the region of the occipital lobe (Verkhlyutov, 1999), which we reproduced in simulation (Fig. 2C).

Thus we can conclude that the propagation of radial waves from certain epicenters can describe the experimental data. In our case, as it can be seen from the table, 8 epicenters for the visual cortex and 1 for the precentral gyrus fulfill this task. We are also able to eliminate all asymmetric cases that give rotational, transverse and diagonal dynamic patterns on the surface of the scalp. In our case, we can conclude that the most likely symmetrical epicenters for fronto-occipital TW on the scalp are located in the visual cortex of both hemispheres in the region of the calcarine and parieto-occipital sulcus. The same dynamics is possible for symmetric epicenters located in the precentral gyrus near the interhemispheric sulcus.

Occipito-frontal TW also can have epicenters in the visual cortex. For other locations of the epicenter, the dynamics of TW has bifurcated extrema of the equal sign (see Table & Supplementary materials).

All the TW trajectories on the scalp are closed, similar to those that were obtained in a work of Hindriks et al. confirming our assumptions about the intracortical nature of electroencephalographic TWs (Hindriks et al., 2014). Similar results were also obtained in a work on 27 subjects, where the authors however did not make assumptions about the nature of the EEG of TW (Manjarrez et al., 2007).

According to our model, the dynamics of the electric field on the surface of the scalp (EEG) for different epicenters should be clearly distinguishable. However, in some cases problems may arise in their identification due to the similarity of the dynamic patterns, as indicated by the high correlation level for some pairs (Fig. 5).

The very first task that can be solved using our method is the localization of the epicenter of the epileptic focus basing on ECoG data (Hindriks, 2016). Two types of electrode matrices - macroelectrode and microelectrode are considered for solving this problem. The results obtained in this case show that TW can be recorded throughout the convectional surface of the brain (Zhang, 2017), but not within the sulci. Therefore, it is difficult to say what amount of data from macroelectrode matrices might arise from volumetric currents, and what amount is related to local field potential (LFP), because these data do not take into account the activity inside the sulci. On the other hand, microelectrode matrices register fragments of radial waves in the parietal, temporal and occipital areas (Martinet, 2017).

Our modeling can fill the gap between the macro and micro scales of leads by recording the potential distribution fragments in ECoG and LFP for microelectrode matrices and comparing them with simulation data.

The outlined considerations allow to building the intuition and lead us into the direction of development of a novel method for solving the inverse problem in EEG and MEG based in the propagating wave *prior*. To properly do this task we will need to formulate a specific functional whose maximization will be equivalent to solving this problem. This will also require the use of individual head geometry and exploiting the advanced techniques for solving the forward problem.

## Methods

TW were modeled in the form of concentric waves propagating over a complex folded surface according to the algorithm developed by authors. The algorithm is implemented as a package of procedures in the Matlab environment which aim to model the electrical potential on a complex folded surface mimicking the human cerebral cortex as a result of the wave excitation process of electrical activity propagating from a single epicenter.

The procedure package consists of the following functions: meshm_dist, meshm_wave, meshm_dipl, meshm_pot (Appendix). Two matrices define triangulated surfaces: Vertices of size 3xN with coordinates of surface nodes and Faces of size 3xL with numbers of three nodes forming L triangles of the surface. All metric values for calculations are specified in mm. The functions work in the following order: 1) the localization of the epicenter as a node of a triangulated surface, 2) the calculation of the geodetic distances to all other nodes of the surface, 3) the calculation of the amplitudes A at each node for the propagating wave, according to the function f(xj,t) basing on the distance from the epicenter xj at each moment time t, 4) calculation of dipole parameters, 5) calculation of potentials.

The epicenter of TW was specified as the target node of the triangulated surface. The user can specify the node directly or the nearest node is determined to a point with user-defined coordinates. The operation is part of the meshm_dist function, which takes a vector with three Cartesian coordinates of the starting point and an array of Vertices, and returns the index of the starting node.

To calculate the geodetic distance from the selected node to all other nodes of the triangulated surface, the latter is represented as a graph, and the graph adjacency matrix is calculated using the Brainstorm toolbox function (Tadel et al., 2011) - tess_vertconn. This function takes two arguments: the vertices and faces of the triangulated surface. As a result, tess_vertconn returns the adjacency matrix of the nodes of the triangulated surface. The adjacency matrix is used for breadth-first search across the graph. In the first iteration, the matrix is multiplied by the vector x = [0 0 0 … 1 … 0 0], where 1 corresponds to the index of the initial node, resulting in the vector y = [0 0 0 … 1 … 0 … 1 … 0], where 1 corresponds to the indexes of nodes adjacent to the first, i.e. its first level of adjacency. The distance between the initial node and its first level of adjacency is calculated. At the next iteration, the adjacency matrix is multiplied by the vector y and the second adjacency level for the starting node is calculated.

The distance to the nodes of the second level of adjacency is calculated by adding the distances to the connected nodes of the first level of adjacency and the distances between the nodes of the first level of adjacency and the initial node. As a result of each iteration, the vector of distances from the initial node to the already passed nodes and a vector with the indexes of these nodes are stored. The procedure is performed by the meshm_dist function, which takes the following arguments: a structure type variable with Faces and Vertices fields specifying the triangulated surface, and a second variable specifying the index of the starting node, or a vector 1x3 specifying the Cartesian coordinates of the starting point. As a result, the function returns a 1xN vector, where each node number corresponds to the geodetic distance to it from the node specified by the user.

Calculation of the amplitudes of the electrical activity wave that propagates along the surface of the brain from the initial node is performed by the function meshm_wave. A 1xN size vector with distances from the initial node to all nodes of the surface, the variable specifying the maximum propagation distance of the wave, the wavelength λ, the number of time readings M, the frequency of the wave oscillations ω in Hz, and the sampling rate (SR) in Hz are taken as arguments. The amplitude was calculated by the formula 1,

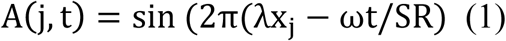

where A (j, t) is the amplitude of the wave at the j-th node point, n is the node index, t is the time moment, and xj is the distance to the j-th node. As a result, the function returns the amplitude matrix NxM for N nodes and M time samples.

The parameters of the elementary dipoles vectors located at the nodes of the surface are calculated by the function meshm_dipl. The function takes a structure type variable with Vertices and Faces fields of the triangulated surface and an NxM amplitude matrix for N nodes and M time samples as arguments. Using the functions of the Brainstrorm software package (http://neuroimage.usc.edu/brainstorm/) tess_normals, the values of unit vectors normal to the surface in each node are calculated. This function takes the Vertices and Faces matrices and returns an array of 3 × N with three Cartesian coordinates of the normal vectors for each of the N nodes. Then, in mesm_dipl, the lengths of the surface normal vectors along which the propagation takes place are multiplied with the amplitudes of the propagating wave.

In addition, the coordinates of the equivalent dipole are calculated as the vector sum of elementary dipoles. Next, the coordinates of the origin of the equivalent dipole are calculated. To do this, cosines are first calculated between each elementary dipole and the equivalent dipole, and then we obtain the radius vectors drawn to the beginnings of elementary dipoles, projected using these cosines. The values of the projections are summed, returning the coordinates of the origin of the dipole. The meshm_dipl function returns a structure type variable with Loc fields of 3xM coordinates of the location of the equivalent dipole, Amp with the size of 3xM amplitudes of the equivalent dipole, and elem with similar Loc and Amp fields of 3xNxM coordinates specifying the position of the elementary dipoles.

Calculation of the electric field on the surface of the head or at selected points simulating the electrodes of the EEG is performed by the function meshm_pot. The function takes the surface of the cerebral cortex, the inner and outer surface of the skull, and the surface of the head, the structure-type variable that specifies the location of the sensors, the vector that specifies the indexing of the sensors, the structure-type variable that specifies the coordinates of the elementary dipoles or the total equivalent dipole as the arguments. The variable specifying the location of the sensors is a structure variable of size 1xK with Name fields, and Loc. Name is the name of the electrode, for example Cz. Loc - location coordinates of electrodes. The method of calculation is given by a string and can be ‘elem’ or ‘equiv’. The first method ‘elem’ calculates the electric field basing on the elementary dipoles, and the second ‘equiv’ basing on the equivalent dipole. Using the bst_openmeeg function of the Brainstorm software package, gain matrix Kx3N (K is the number of channels) is calculated for the first time point. Next, for each time point, the value of the electric or magnetic field potential on the electrodes is calculated by multiplication of elementary dipoles vectors with the gain matrix and written into a matrix of the size KxM, which is returned by a function meshm_pot. In the case of the ‘equiv’ method, the gain matrix is calculated for each time point and multiplied by the coordinates of the equivalent dipole corresponding to this time point.

An equivalent dipole was calculated from a set of dipoles perpendicular to the model surface. The algorithm allowed to calculate the electrical potentials on the model surface of the scalp at the locations of the 129 electrodes of the Geodesics system (Electrical Geodesics Inc., USA), including the 129th reference electrode Cz. In order to do this, the boundary element method was implemented in the OpenMEEG software (Kybic et al., 2005; Gramfort et al., 2010). As a result a 100-msec model EEG was generated, reproduced by the Brainstorm program (Tadel et al., 2011), and the interpolated dynamical distribution of the electric field on the scalp was mapped. At the same time, EEG sources were localized using the standard wMNE method implemented in the Brainstorm toolbox (Tadel et al., 2011).

In the computer model of the wave propagating along the complex folded structure of the human cerebral cortex, the current density was set by a unit vector V which was collinear to the normals of the triangulated surface and varied sinusoidally from −1 to +1 depending on the geodetic distance from the epicenter. To define the current density in nA / mm2, a special procedure was used to calculate the total area S occupied by the propagating wave process. Then the number of normals Nv located on this area and the area per one normal Sv = S / Nv were calculated, and then we estimated the current at the normal CDv = V * CD / Sv, where CD is the current density in A / m2 (nA / mm2). The current density in different models was set to vary from 50 to 250 nA / mm2, according to measurements from the visual evoked response registration from the monkey cortex (Hamalainen, et al., 1993). Basing on the model of the cortical surface ICBM152 from the Brainstorm toolbox (http://neuroimage.usc.edu/brainstorm/) the distribution of vectors and current density values estimated on them was computed in a geodesic radius of 4 cm from the epicenter for 50 time points, i.e. every 2 ms, which at a sampling rate of 500 Hz was 100 ms. Thus a complete unit wave with a frequency of 10 Hz was formed, propagating at a velocity of 0.2 m / s (Verkhlyutov, 1996).

We modeled 28 wave patterns per 100 milliseconds of the model EEG and the dipole distributions in the cortex were calculated. According to the described procedure we calculated an equivalent dipole and simulated EEGs.

The estimated EEG temporal patterns in the form of three-dimensional matrices were compared with each other using the corr2 procedure from the Matlab standard library. We calculated mean, minimal, maximal values (except a value of 1) and first and third quartiles of mutual correlation of the simulated EEG from radial waves propagating from 28 epicenters on the ICBN152 cortical surface model (Fig. 1).

## Acknowledgements

V.V. acknowledges support from RFBR Grant №17–04–02211a. Raw data for this paper are available at http://braintw.org/. The authors have no competing financial interests.

## Appendix

~~~
% Geodesic distances
% mesh_distance-distances to all vertices
% mesh – surface
% Face – vertex number or coordinates of vertex
function [mesh_distance] = **meshm_dist**(mesh,Face)
       tic
       Faces=mesh.Faces;
       Vertices=mesh.Vertices;
       VertConn=tess_vertconn(Vertices,Faces);
       % VertNormals = tess_normals(Vertices, Faces, VertConn);
       nv = size(Vertices,1);
       face=find_nearest_face(Face,Vertices,nv);
       mesh_distance = distance_count( face, Vertices, VertConn, nv);
       toc
end
% mesh_distance-distances to all vertices
% face-start vertex
% nv-number of vertices
function [mesh_distance] = distance_count(face, Vertices, VertConn, nv)
              VertConnGrow = zeros(1,nv);
              VertConnGrow(face)=1;
              mesh_distance=zeros(1,nv);
              Iter=0;
              while 1
                       Iter=Iter+1;
                       VertConnPrev = VertConnGrow;
                       VertConnGrow = double(VertConnGrow * VertConn > 0);
                       if Iter>1
                       VertConnGrow(find(mesh_distance>0))=0;
                       end
                       VertConnGrow(face)=0;
                       vind = find(VertConnGrow - VertConnPrev > 0);
                       for i=1:length(vind)
                             mesh_distance(vind(i))=0;
                             vold=find(VertConnPrev > 0);
                             if Iter>1
                                PrevConn=find(VertConn(vold,vind(i)));
                                mesh_distance(vind(i))=sqrt((Vertices(vold(PrevConn(1)),1)-…
                                Vertices(vind(i),1)).^2+(Vertices(vold(PrevConn(1)),2)-…
                                Vertices(vind(i),2)).^2+(Vertices(vold(PrevConn(1)),3)-…
                                Vertices(vind(i),3)).^2)+mesh_distance(vold(PrevConn(1)));
                             continue;
                             else
                               mesh_distance(vind(i))=sqrt((Vertices(face,1)-…
                               Vertices(vind(i),1)).^2 + …
                               (Vertices(face,2) -…
                               Vertices(vind(i),2)).^2 + …
                               (Vertices(face,3) - Vertices(vind(i),3)).^2);
                            end
                      end
                      if Iter>2
                         if numel(find(mesh_distance)>0)==…
                            ndist||numel(find(mesh_distance)>0)==nv-1
                             break;
                            end
                      end
                     if Iter>1
                        ndist=numel(find(mesh_distance)>0);
                    end
         end
end
function [face]=find_nearest_face(Face,Vertices,nv)
if size(Face,2)==3
      parfor i=1:nv
            d(i)=sqrt((Vertices(i,1) - Face(1)).^2 + …
                  (Vertices(i,2) - Face(2)).^2 + (Vertices(i,3) - Face(3)).^2);
      end
      face=find(d==min(d));
      disp(strcat(‘Initial face is #’,num2str(face)));
else
      face=Face;
end
end
% distances array  (mesh_dist)
% max distace       (max_dist)
% length wave       (l_wave)
% number steps      (N_step)
% wave frequency  (w),
% sampling rate    (SR)
function [Amp] = **meshm_wave**(mesh_dist, max_dist, l_wave, N_step, w, SR)
Amp=[];
md=(mesh_dist<max_dist)-(mesh_dist==0)-(mesh_dist==Inf);
for i=1:N_step
       Amp(:,i)=sin(2*pi*mesh_dist/l_wave-2*pi*i*w/SR).*md;
end
end
function [ dipe ] = **meshm_dipl**( mesh, Amp)
Vertices=mesh.Vertices;
Faces=mesh.Faces;
VertNormals = mesh.VertNormals;
N_step=size(Amp,2);
md=(Amp(:,2)~=0);
md3=repmat(md,1,3);
for i=1:N_step
      for k=1:3
            DipAmplitude(i,k)=…
                  sum(VertNormals(:,k).*(Amp(:,i)))/numel(VertNormals(:,1));
      end            Dip_proj(:,i)=sum(VertNormals.*md3.*…
      repmat(DipAmplitude(i,:),size(VertNormals,1),1),2);
      Dip_proj((Dip_proj(:,i))<0,i)=0;
DipLoc(i,:)=sum(Vertices.*repmat(Dip_proj(:,i),1,3),1)/sum(Dip_proj(:,i));
      DipAmplitude(i,:)=DipAmplitude(i,:)+DipLoc(i,:);
end
dipe.Loc=DipLoc;
dipe.Amp=DipAmplitude;
end
function [Rec]=**meschm_pot**( cortexfile,inner,head,outer,channel,iEeg,dipe, elem)
% Creating OPTIONS structure for bst_openmeeg
            OPTIONS.Comment=‘’;
            OPTIONS.HeadModelFile=head;
            OPTIONS.HeadModelType=‘surface’;
            OPTIONS.Channel=channel;
            OPTIONS.MEGMethod=‘’;
            OPTIONS.EEGMethod=‘openmeeg’;
            OPTIONS.ECOGMethod=‘’;
            OPTIONS.SEEGMethod=‘’;
            OPTIONS.CortexFile=cortexfile;
            OPTIONS.InnerSkullFile=inner;
            OPTIONS.BemFiles={head, outer, inner};
            OPTIONS.BemNames={‘Scalp’ ‘Skull’ ‘Brain’};
            OPTIONS.BemCond=[0.33 0.0041 0.33];
            OPTIONS.iEeg=iEeg;
            OPTIONS.BemSelect=[1 1 1];
            OPTIONS.isAdjoint=0;
            OPTIONS.isAdaptative=1;
            OPTIONS.isSplit=0;
            %Checking for elem option, if ‘equiv’ then only one dipole is used for
            %forward modelling (dipe.Loc - location of dipole, dipe.Amp - the
            %location of an end of dipole. Dipole vector is dipe.Amp-dipe.Loc).
            % if ‘elem’ then dipoles are used which are placed in the vertices of
the
            % mesh (dipe.elem - elemental dipoles. only
            % then iterating for time points (i)
            if strcmp(elem,‘equiv’)
            for i=1:size(dipe.Loc,1)
                    % location of the dipole and its orientation (via unit vector)
                    OPTIONS.GridLoc=dipe.Loc(i,:);
                    OPTIONS.GridOrient=(dipe.Amp(i,:)-…
                    dipe.Loc(i,:))/norm(dipe.Amp(i,:)-dipe.Loc(i,:));
                    G=bst_openmeeg(OPTIONS);
                    Rec(:,i)=G*(dipe.Amp(i,:)-dipe.Loc(i,:))’;
            end
            elseif strcmp(elem,’elem’)
                    OPTIONS.GridLoc=squeeze(dipe.elem.Loc(1,:,:));
                    OPTIONS.GridOrient=…
                    squeeze((dipe.elem.Amp(1,:,:))/norm(squeeze(dipe.elem.Amp(1,:,:))));
                    G=bst_openmeeg(OPTIONS);
                    for i=1:size(dipe.Loc,1)
                            Rec(:,i)=G*reshape(((dipe.elem.Loc(i,:,:)-…
                            dipe.elem.Amp(i,:,:))),size(G,2), 1);
                    end
            end
end
~~~

## Supplementary materials

We recommend looping the animation when viewing.

### Movie 1

**Modeling the EEG and equivalent dipole from a radial wave in the visual cortex**. Modeled EEG for 100 ms. Electric field distribution on the “scalp” (the occipital-frontal propagation) from a wave in from two symmetric waves in both hemispheres with an epicenter at the vertex 03.008488 of the cortical surface model ICBN152 (the second wave is mirror-wise with respect to the sagittal plane). The equivalent dipole rotates in the sagittal plane at the time indicated by the cursor in EEG window. The current density distribution from radial wave on the unsmoothed and smoothened cortical surface model.

### Movie 2

**Modeling the EEG and equivalent dipole from a radial wave in the visual cortex**. Modeled EEG for 100 ms. Electric field distribution on the “scalp” (the counter-clockwise rotation) from a wave in left hemisphere with an epicenter at the vertex 04.18914 of the cortical surface model ICBN152. The equivalent dipole rotates in the axial plane (top view) at the time indicated by the cursor in EEG window. The current density distribution from radial wave on the unsmoothed and smoothened cortical surface model.

### Movie 3

**Modeling the EEG and equivalent dipole from a radial wave in the sensorimotor cortex**. Modeled EEG for 100 ms. Electric field distribution on the “scalp” from a wave in left hemisphere with an epicenter at the vertex 19.071133 of the cortical surface model ICBN152. The current density distribution from radial wave on the cortical surface model. The equivalent dipole is precesses within the Rolando fissure.

*Movies on the site* http://braintw.org

main->Cortex TW Simulation->Movie->

170226movCORsim (Cortex TW simulation from 28 epicenters), 170330movEEGsim (EEG TW simulation for 28 epicenters, left and both hemispheres).

